# Tickle contagion in the somatosensory cortex

**DOI:** 10.1101/2021.10.22.465426

**Authors:** Lena V. Kaufmann, Michael Brecht, Shimpei Ishiyama

## Abstract

The cellular mechanisms of emotional contagion are unknown. We investigated tickle contagion and the underlying neuronal representations in rats. We recorded trunk somatosensory cortex activity of observer rats while they received tickling, audio-visual playback of tickling footage, and while they witnessed tickling of demonstrator rats. Observers vocalized, and showed “Freudensprünge” (“joy jumps”) during witnessing live tickling, while they showed little behavioral responses to playbacks. A fraction of trunk somatosensory neurons responded to both direct and witnessed tickling in action-specific manner. The correlation between direct and witnessed tickling responses increased towards deeper cortical layers. Tickle-mirror neurons but not non-mirror neurons discharged prior to and during vocalizations and hence might drive contagious ‘laughter’. We conclude that trunk somatosensory cortex represents mirrored ticklishness.

## Introduction

The capacity of an individual share the feelings other individuals [1] plays a critical role in human social interactions [2]. Possible mechanisms for this phenomenon have for a long time been a subject of discussions in various disciplines [3]. From an evolutionary point of view, emotional contagion, “feeling into” one’s conspecifics, can be a vital ability, not only for humans but for all social species [4-6]. Emotional contagion as a way to gain rapid emotional connectedness has been proposed to have its origins in parental behavior and is thought to be the root of empathy and a precursor of prosocial behavior [7]. There is a growing field of research on rodent empathy, but until now these studies have mainly been focusing on negative emotions [8]. Empathy is a rather ambiguous term and here the focus will lie on its basal form of emotional contagion as “primal empathy” [1] or “affective empathy” [5] rather than higher, cognitive embodiments of empathy.

In addition to the higher order visual system, the mirror neurons have been suggested to be involved in understanding others actions [9]. Since their first discovery in macaque ventral premoter area as neurons responding not only to the execution but also to the observation of specific motor actions [10, 11], mirror neurons have been described in various brain regions in macaques [12-17] and humans [18]. Recent studies found roles of mouse anterior cingulate cortex-to-nucleus accumbens projections in the social transfer of pain [19], the mouse suprachiasmatic nucleus in itch contagion [20], and the rat anterior cingulate cortex containing cells that respond to experienced and witnessed pain – “emotional mirror neurons” [21]. Interestingly, somatosensory regions have also been described to play a role in the visual recognition of emotion, and the reactivation of internal somatosensory representations is necessary to simulate the emotional state of another [22-25].

Despite many behavioral observations of the contagion of ticklish laughter [26], so far there are few studies investigating what happens in the brain when the mere observation of others being tickled and laughing instantly makes us giggle as well. With help of advances that provided evidence of ticklishness in rats [27] and the neural correlates of ticklishness [28], we investigated whether the rats’ behavioral response to tickling is contagious and whether mere observation of a demonstrator rat being tickled is sufficient to induce a measurable positive emotional state in an observer rat. We describe behavioral observations suggestive of “laughter contagion” or more generally contagion of positively excited behavior in rats. We show here that there is indeed a subset of “tickling mirror-like neurons” in the trunk region of the somatosensory cortex.

## Results

Observer (= subject) rats were placed in a compartment of a box separated from a demonstrator compartment (Figure 1A). The observer rats received tickling on the dorsal and ventral trunk by the experimenter, and air tickling (experimenter made a tickling motion in the demonstrator compartment). The observer rats then received audio, visual and audio-visual playback of rat tickling footage (Figure 1A right). Finally, a demonstrator rat was introduced and tickled (witnessed tickling, Figure 1A, left). The experimental paradigm is illustrated in Figure 1B. The subject rats seemed to pay attention to the stimuli, quantified as fractions of time where the head was directed to the demonstrator compartment (air tickling: 80 ± 7%, *n* = 4 recordings; audio/visual playback: 65 ± 5%, *n* = 4; visual playback: 62 ± 10%, *n* = 3; demonstrator introduction: 79 ± 7%, *n* = 4; demonstrator tickled: 85 ± 8%, *n* = 4). Differences in sound intensity of ultrasonic vocalizations (USVs) captured by four microphones, combined with positions of the animals relative to the microphones, allowed us to assign the emitter of most (90.8%) calls. As shown previously [28, 29], the observer rats strongly responded to tickling with USVs (Figure 1C, top) and *Freudensprüngen* (“joy jumps”; Figure 1C bottom). Audio and/or visual playback of tickling footage had little effect on the observers’ USVs and jumps (Figure 1D&E). Interestingly, however, the observer rats vocalized and jumped when the demonstrator rats were tickled (Figure 1D&E; Movie S1).

**Figure 1.**
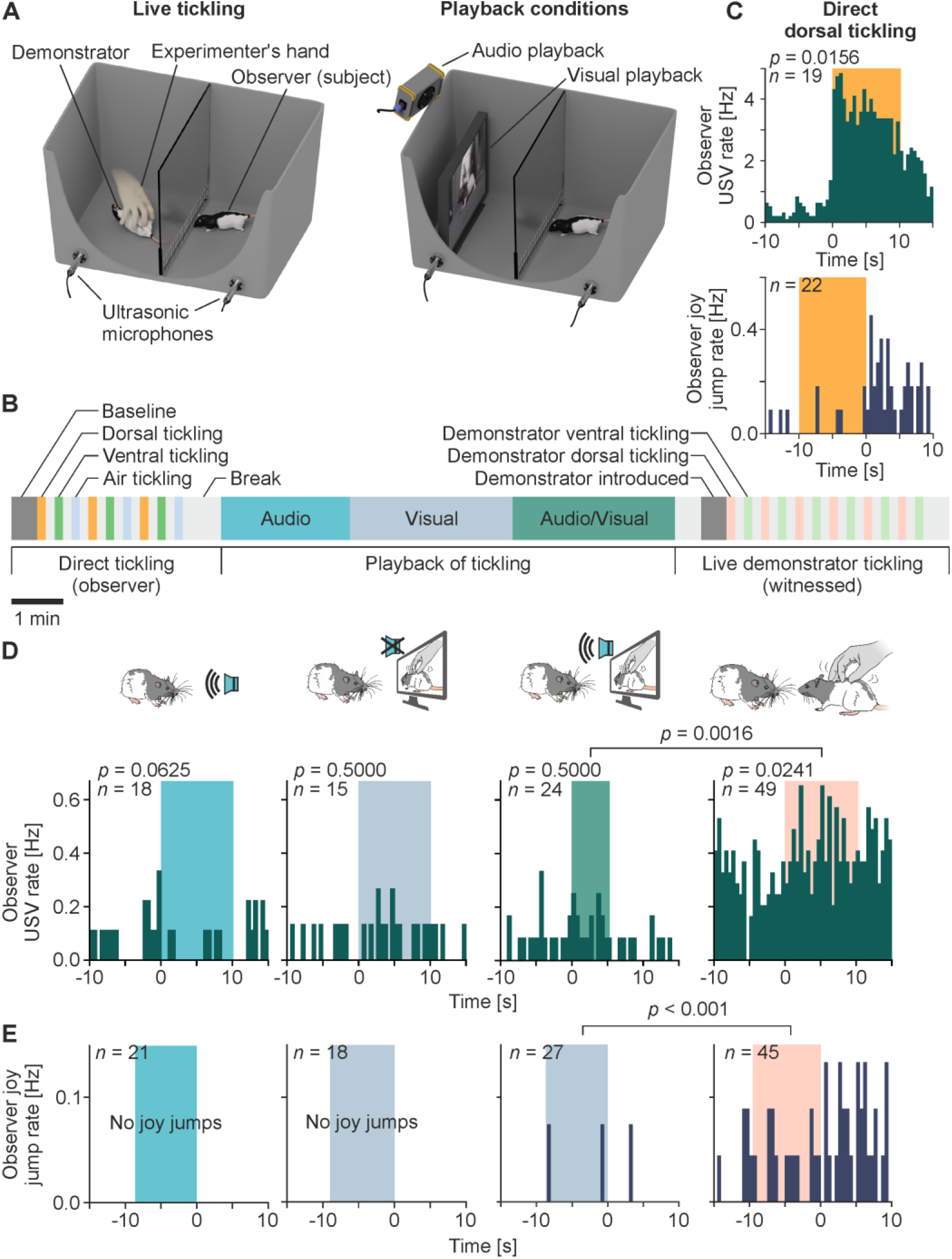
Experimental setup and behavioral response to witnessed tickling. A. Experimental setup for live demonstration (left) and audio/visual playback (right) of tickling. The box was separated by a transparent acrylic partition with a steel mesh at the bottom. B. Timeline of the experimental paradigm. Color boxes indicate different event phases. Order of playback stimuli (audio; visual; audio/visual) was randomized in each recording. C. Peri-stimulus time histograms (PSTHs) of ultrasonic vocalization (USV) rate (top), and Freudensprünge (“joy jumps”, bottom) rate aligned to the onset and offset of direct dorsal tickling, respectively (bin width = 0.5 s). Width of the color boxes indicate median duration of tickling. *n* refers to number of events (USVs: 19 tickling events, 4 recordings from 4 rats; jumps: 22 tickling events 7 recordings from 4 rats). *p*-value: signed-rank test for before vs. after the event onset. D. Top, schematic illustrations of (from left) audio playback, visual playback, audio/visual playback, and witnessed dorsal tickling events. Bottom, PSTHs of observer USV rate, aligned to the onset of events depicted above (bin width = 0.5 s). Width of the color boxes indicate median duration of events. *n* refers to number of events (audio: 18 events, 6 recordings, 4 rats; visual: 15 events, 5 recordings, 3 rats; audio/visual: 24 events, 8 recordings, 4 rats; witnessed: 49 events, 8 recordings, 4 rats). *p*-value on each chart: signed-rank test for before vs. after the event onset. Comparisons of USV rates during audio/visual dorsal tickling vs. witnessed dorsal tickling: rank-sum test. E. Observer Freudensprünge rates aligned to the offset of the events same as (E). *n* refers to number of events (audio: 21 events, 7 recordings, 4 rats; visual: 18 events, 6 recordings, 3 rats; audio/visual: 27 events, 7 recordings, 4 rats; witnessed: 45 events, 7 recordings, 4 rats). Comparison between audio/visual and witnessed dorsal tickling was calculated with frequency of jumps from phase onset to 10 s after phase offset (rank-sum test).

We recorded extracellular unit activity in the observers’ trunk somatosensory cortex. A representative unit in layer 5b strongly responded to direct dorsal tickling, air tickling and witnessed demonstrator dorsal tickling (Figure 2A). We calculated a response index (RI, see methods) for each unit and for each event. Of all 568 recorded units, 76%, 48% and 38% units were activated (RI > 0.1) by direct dorsal tickling, air tickling and witnessed tickling, respectively (Figure 2B). We defined units fulfilling both RI_witnessed_dorsal_ > 0.1 and RI_observer_dorsal_ > 0.1 as ‘tickling mirror units’, and 32% of all recorded units appeared to be mirror units. In population, units that were activated by direct dorsal tickling had RIs for air tickling significantly shifted towards positive values (Figure 2C, top). Units that were inhibited by direct dorsal tickling did not have RIs for air tickling shifted from zero (Figure 2C, bottom). RIs for air tickling in units that were activated by direct dorsal tickling were significantly larger than those in units that were inhibited by direct dorsal tickling (Figure 2C). The same trait was found also in the response to witnessed tickling (Figure 2D), suggesting a neuronal link between ticklishness, witnessed tickling hand and tickled conspecifics.

**Figure 2.**
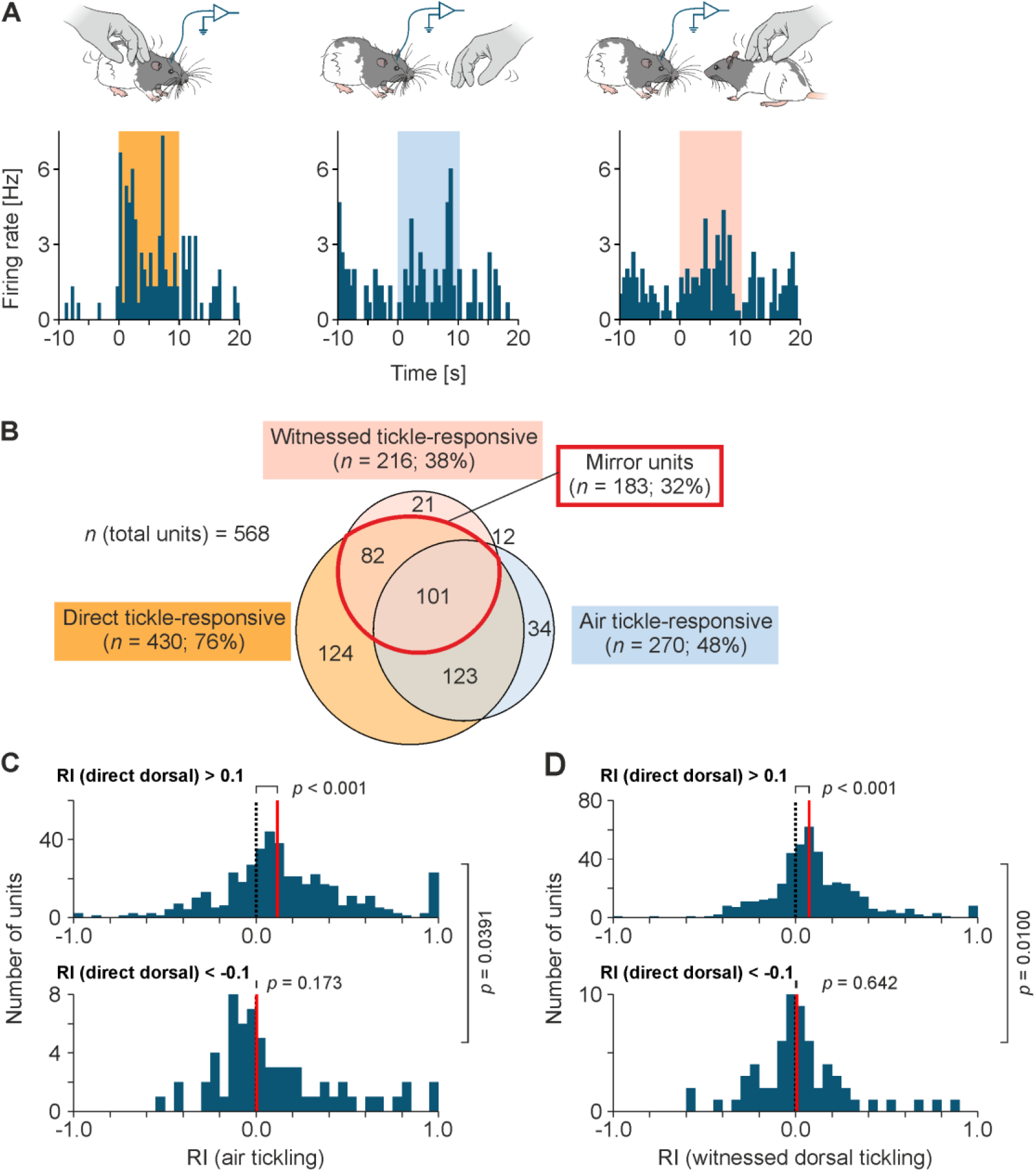
Somatosensory cortical neurons respond to tickling, air tickling, witnessed tickling and such responses are related. A. Top, schematic illustrations of neuronal recording during tickling (left), air tickling (middle, hand with tickling motion without touching), and witnessed tickling of demonstrator (right). Bottom, peri-stimulus time histograms of firing rate of a representative unit. Data are binned to 0.5 s. Width of the color boxes indicate median duration of events. B. Venn diagram of response pattern of the total units recorded (*n* = 568 units). Response pattern was calculated as response indices (RIs, see methods) > 0.1 in observer dorsal tickling (Tickle-responsive), witnessed dorsal tickling of demonstrator (Witnessed tickle-responsive), and air tickle (Hand-responsive). Mirror units (red) were defined as units responding to both tickling and witnessed tickling. C. Top, histogram of RIs for air tickling in units where RIs for observer tickling were larger than 0.1. The *p*-value indicates one-sample *t*-test against 0. Bottom, same as top but in units where RIs for observer tickling were smaller than −0.1. The *p*-value at right indicates rank-sum test between the two populations. D. Same as C but for RIs for witnessed dorsal tickling.

We next analyzed the neuronal response to direct tickling vs. witnessed tickling between cortical layers (Figure 3). Cortical layers were assigned according to cytochrome C oxidase staining (Figure 3A). Interestingly, deep-layer (5b; 6) but not superficial-layer (2/3; 4; 5a) units showed a significant correlation between RIs for direct and witnessed tickling (Figure 3B&D). The same trait was found for direct tickling vs. air tickling (Figure S1). On contrary, fraction of mirror units was the smallest in layer 6 (Figure 3C).

**Figure 3.**
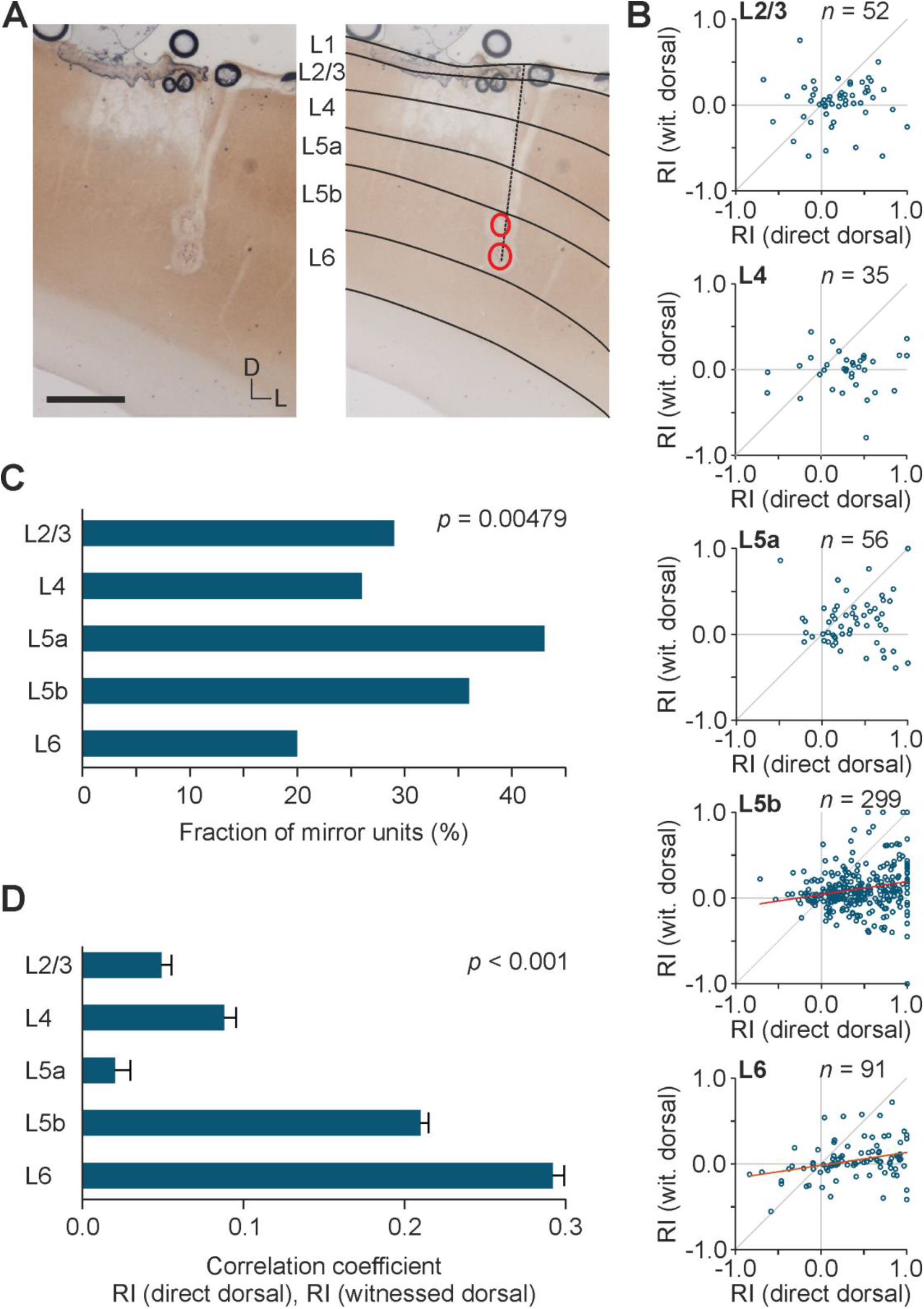
Tickle response predicts witnessed tickle response in deep-layer trunk cortex. A. Representative histological analysis for layer assignment. Left, a coronal brain section stained for cytochrome C oxidase (scalebar = 0.5 mm). Right, assignment of cortical layers (black lines). Dashed line: tetrode track. Red circles: lesions. B. Scatter plots show response indices (RI) for dorsal tickling (dorsal) vs. witnessed dorsal tickling of demonstrator (wit. dorsal) in layers 2/3, 4, 5a, 5b and 6. Grey diagonal lines are unity line. Red lines are linear fit in case of significant correlation (Pearson’s). *n*: number of units. C. Fraction of mirror units i.e. RI (witnessed dorsal) > 0.1 & RI (dorsal) > 0.1 in each layer. *p*-value: chi square test. D. Pearson’s correlation coefficient *r* calculated for 1000 times with randomly chosen 21 units, i.e. 60% of layer 4 sample size, in each layer. Mean ± SEM; *p*-value: Friedman test.

Whereas responses to direct and witnessed tickling were correlated, there were no systematic correlations between responses to direct tickling and playback of tickling footage across layers (Figures S2 & S3).

We wondered whether the neuronal response to air tickling and witnessed tickling has any action or contextual specificity. Most of our recorded units responded more strongly to dorsal touch than ventral touch, resulting in a stronger response to flip (the experimenter grabbed the dorsal trunk and restrained on the back) than to ventral tickling (representative unit, Figure S4A, left). The stronger response to flip than ventral tickling was readily obvious also during witnessed tickling in the same unit (Figure S4A, middle). Witnessed dorsal tickling, on the other hand, did not evoke a strong pre-tickling response as seen in witnessed flip followed by witnessed ventral tickling (Figure S4A, right). Layer 5b population firing rates were higher during witnessed demonstrator flip than during witnessed demonstrator ventral tickling (Figure S4B). These results suggest that trunk somatosensory cortex responds to witnessed tickling in an action specific manner, mirroring the tickling experienced by the demonstrator.

We previously showed that the trunk somatosensory neurons increase their firing rate upon USV emission [28]. We here compared the neuronal response to witnessed and own USV emission in mirror units and non-mirror units (Figure 4). Mirror units did not respond to demonstrator calls (Figure 4A & B, left) but strongly responded to own calls (Figure 4A & B, right). Interestingly, non-mirror units responded to neither demonstrator nor own calls (Figure 4C&D).

**Figure 4.**
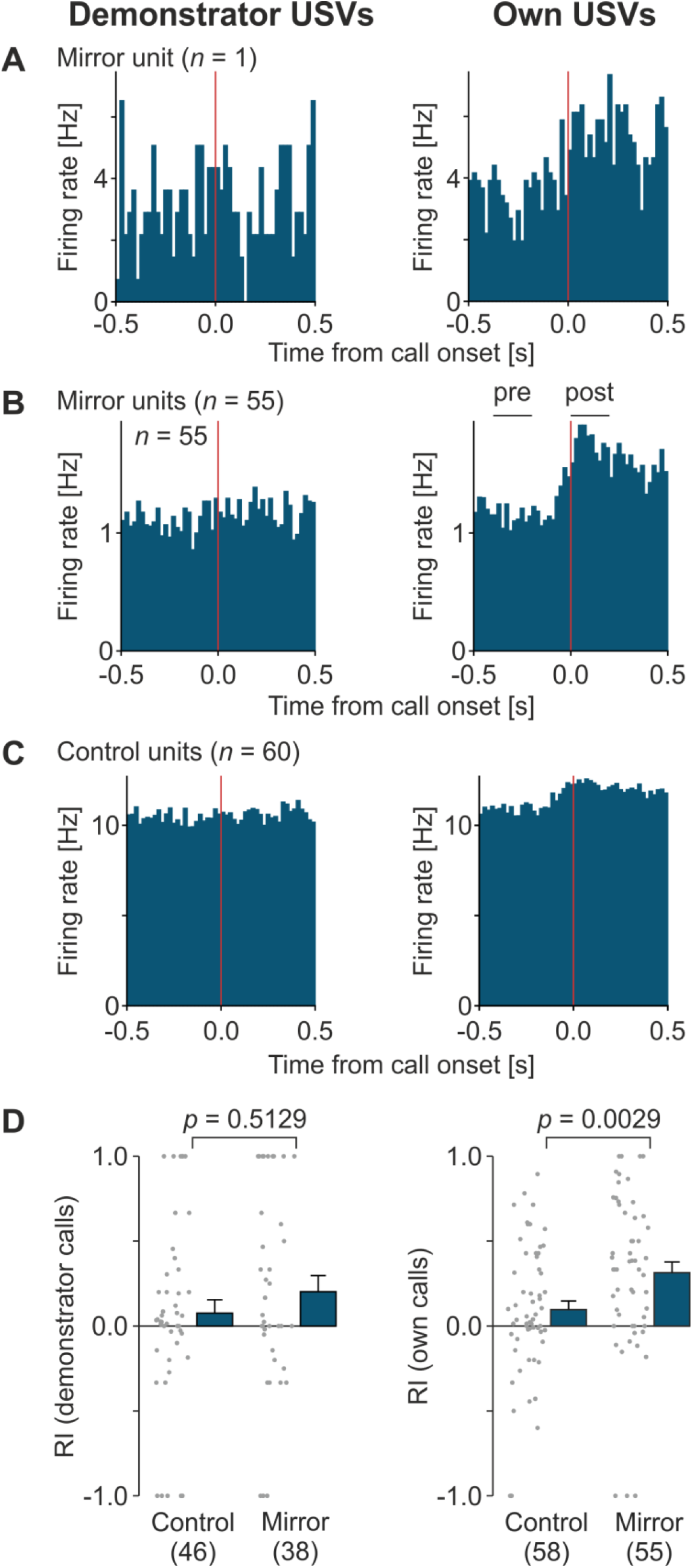
Mirror-units discharge prior to and during own vocalizations. A. Representative peristimulus time histogram (PSTH) of a mirror unit (responding to both tickling and witnessed tickling measured as response index (RI) > 0.1, see methods) in layer 5b trunk somatosensory cortex, aligned to the onset of demonstrator ultrasonic vocalizations (USVs, in break phases with no previous USV onset within 0.5 ms, left) and own USVs (right). Data are binned to 20 ms. B. Population PSTH of 55 mirror units aligned to demonstrator USV onset (left) and own USV onset (right). ‘pre’ and ‘post’ on the right panel indicate time ranges to calculate response indices in (D). C. Same as (B) but for control units (responding to tickling i.e. RI > 0.1, but not responding to witnessed tickling i.e. −0.1 < RI < 0.1). D. Firing rate response indices (RIs) for demonstrator USVs (left) and own USVs (right). RIs were calculated as (FR_post_ – FR_pre_) / (FR_post_ + FR_pre_), where FR_pre_ is average firing rate in [−0.4, −0.2] s to USV onset, and FR_post_ is average firing rate in [0.0, 0.2] s to USV onset, as indicated in (B). Number of units are indicated at the bottom. Comparisons between control and mirror units were performed by Mann-Whitney rank-sum test.

## Discussion

We verified our previous results [28] that rats show a neuronal and behavioral response to tickling. Rats showed an apparent interest in the conspecifics being tickled. Fifty-kHz USVs are reliable measures of positive emotions in rats [30] and *Freudensprünge* are described as a behavior that is seen in joyful rats [28, 31]. We observed such excited behavior in the observer rat witnessing the demonstrator rat being tickled, pointing to emotional contagion of play behavior. This kind of playfulness contagion has been observed in e.g. ravens [32] and keas [33]. Contagion of laughter has been observed in humans [34] and chimpanzees [35]. Several studies on rats showed an increase in play by introduction of a more playful individual [36, 37] and tickling-induced emotional contagion from a tickled rat to their cage mate [38]. Devocalized pairs of rats show reduced play frequency [39]. Artificially deafened but not blinded rats [40] play less than their conspecifics [41]. We found that the observer rat showed little behavioral response to audio and/or visual playback of tickling footage (Figure 1D&E), but more to the demonstrator rat being tickled, suggesting vicarious playfulness. Yet, it remains to be tested whether playback or live demonstration of play leads to an increase in play behavior in observer animals.

Next, we found that trunk somatosensory neurons responded to witnessed tickling, and 32% of the units increased their firing rate during both direct and witnessed tickling (‘tickling mirror-units’, Figure 2B). The fraction of the mirror units approximately agrees with the number of the mirror neurons reported in macaque premotor cortex [11]. This is a particularly interesting discovery, since sensory mirror neurons are not well studied and, to our knowledge, this is the first report of sensory mirror-like neurons in the primary somatosensory cortex. Moreover, this neuronal response was associated with positive emotional behavior.

We found higher percentages of tickle-mirror units in layers 5a, 5b than in other layers (Figure 3C). Only deeper layers (layer 5b and 6) showed a significant correlation between direct and witnessed tickling (Figure 3D). Fraction of mirror units was the largest in layer 5a, whereas layer 6 had the smallest fraction of mirror units, as the responses varied from suppression to excitation. Traditionally, the somatosensory cortex is seen as a sensory map with layer 4 being a sensory relay center but recently we proposed a broader capability of the somatosensory system as a body model that is not only receiving and relaying sensory inputs but serves as a representation of the body and body simulation [42]. We previously showed that discharge of layer 4 trunk somatosensory neurons is locked to tickling, whereas layer 5 plays a role in anticipation of tickling [29]. Layer 5 neurons also show strong responses during tickling, and microstimulation of deep layers evokes ultrasonic vocalizations [28]. The role of deep layers of the trunk somatosensory cortex in tickling and mirror-like responses to tickling could indicate that the deep layers are more related to an emotional response.

Many cells that responded to direct tickling were also excited by the air tickling (Figure 2A-C). There are at least two possible explanations for this phenomenon: The timing of the air tickling could have favored a strong response, since it was performed immediately after and before direct tickling bouts (Figure 1B), which evoked the strongest behavioral and neuronal response. A second possibility would be that these cells responded rather to the expectancy of being tickled than the observed tickling act itself. Previous studies also found that somatosensory neurons fire apparently in response to expected tickling during hand chase [28] and slow hand approach [29]. However, the trunk somatosensory neurons responded not only to the hand for anticipation but also to what the hand does: recorded units that were more strongly excited by flip phase than following ventral tickling phase responded to witnessed tickling in the similar manner (Figure S4). Thus, the tickling-mirror units represent the context i.e., action specificity of witnessed tickling, as if the observer rat itself was being tickled.

The tickling mirror-like units showed a very weak if not no response to the audio and/or visual playback of conspecific tickling (Figure S2). There are actually many accounts of modality-specific mirror neurons, at least two studies found auditory mirror neurons in macaques [43, 44] and neuroimaging in humans led to the proposition of a somatotopic auditory mirror system, most of this proposed system, however, appears to be multimodal [45]. Recently, a study revealed modulation of the barrel cortex by the superior colliculus in mice [46]. Regarding the rat ecology as nocturnal and not very visual animals that are prey for a number of hunters and rely more on their hearing and olfaction [47], more cells might respond to auditory stimuli or these responses could be stronger than to other sensory inputs. Anatomical tracing experiments could potentially reveal more about multisensory integration in the somatosensory cortex.

A recent study demonstrated that rats distinguish USV emitter, and they self-administer preferentially playback of 50-kHz USVs emitted by a stranger rat over those emitted by their cage mate [48]. Using playback footage and live demonstration of a stranger rats, therefore, may have been more rewarding stimuli to the observer rats. It is to be noted, however, that familiarity with the demonstrator is crucial in empathetic behavior of the observer in rodents, at least for negative experiences [49-52].

Carrillo et al. [21] showed that there are emotional mirror neurons in the rat anterior cingulate cortex. The central amygdala also appears to be involved in the recognition and emotional contagion of socially signaled danger [53, 54]. It is likely that the response to direct and witnessing tickling that we observed in the somatosensory cortex neurons was strongly influenced by or dependent on the positive emotional valence of the tickling. Interestingly a number of studies report that the recognition of emotion expression in faces [22, 55], and voices [56, 57] depends on the right somatosensory cortex. Furthermore, it has been proposed that the somatosensory cortex plays a role in linking the perception of emotional expressions with subjective sensory experience [23].

Taken together our results suggest that rat “laughter” and ticklishness is contagious. Hence, our research opens new avenues to investigate positive emotions, which are rather understudied in neuroscience. Much like in humans, live performance (i.e. demonstrator tickling) was much more engaging in rats than the various forms of playback (i.e. watching TV). Mapping of others’ body representations on a “body model” in the somatosensory cortex [42], facilitated by emotional valence would be a possible explanation for these findings.

## Acknowledgements

Supported by BCCN Berlin, Humboldt-Universität zu Berlin, Deutsche Forschungsgemeinschaft: 393810148 and the Leibniz Prize. We thank Undine Schneeweiß, Arnold Stern, Maik Kunert, Viktor Bahr, Falk Mielke, Miguel Concha-Miranda, and Andreea Neukirchner. LVK, MB and SI designed experiments. LVK and SI conducted experiments and analyzed data. All authors wrote the manuscript. The authors declare no conflict of interest. All data generated in this study are available upon request.

## Supplementary Materials

Materials and Methods

Fig S1 – S4

Movie S1

## Materials and Methods

### Animals

Seven 3-week-old male Long-Evans rats were acquired from Janvier Labs. Animals were individually housed, maintained with a 12:12 h inverted light/dark circle and allowed *ad libitum* access to food and water. Prior to the experiments, animals were handled for 20 minutes and tickled for 10 minutes [28] individually once a day for 6-8 days. Additionally, the rats were grouped into pairs, and accustomed to the experimental setup as well as each other every day during the training period to avoid any novelty effect later during the experiment. At the end of the training period the rats responding with more excitement and more ultrasonic vocalizations (USVs) to tickling were chosen as the observer rats and the less responsive ones as the demonstrator rats. All experimental procedures were performed according to German guidelines on animal welfare under the supervision of local ethics committees in accordance with the animal experimentation permit (Permit No. G0193/14 and Permit No. G0279/18).

### Experimental setup

The experimental setup consisted of a box (‘tickle box’) lined with black foam rubber, with a plexiglass separation wall in the middle. At the bottom of the separation wall was a mesh wire stripe, allowing the animals to see, hear and smell each other. The experimental environment was kept dark (< 20 lx). Video material was recorded under infrared illumination with two to three cameras: a top view and a side view camera with 30 fps (Imaging Source, Germany), and a side view with 240 fps (H5PRO – modified GoPro Hero 5, Back-bone, Ottawa, Canada). Ultrasonic vocalizations were recorded with four microphones (condenser ultrasound CM16/CMPA, frequency range 10-200 kHz, Avisoft Bioacoustics, Berlin, Germany) at a sampling rate of 250 kHz and with a 16-bit resolution using Avisoft-RECORDER USGH software (Avisoft Bioacoustics, Berlin, Germany). The microphone positioning enabled assigning calls to demonstrator and observer rat, respectively. The setup is illustrated in Figure 1A.

### Experimental paradigm

Rats were tickled as previously described [28]. The experiments started with the observer rat alone, positioned in the left compartment of the tickle box. A baseline phase was followed by three rounds of dorsal tickling, ventral tickling and air tickling (tickling motion) in the right compartment as a control for the presence of the hand. Each phase of tickling lasted for 10 seconds interleaved by 15 second breaks. This was followed by playback of rat tickling footage, which could be only audio, only visual or audio-visual playback. For the latter two a small display was placed in the right compartment facing the observer rat. The order of the playback phases was randomized. For the last part of the experiment, the demonstrator rat was introduced and after a baseline observation there were six rounds of dorsal and ventral tickling of the demonstrator rat i.e. witnessed tickling. The procedure is illustrated in Figure 1B.

### Video analysis

Video material was analyzed using ELAN 5.1 software [58] to label durations of the experimental paradigm phases as well as rat behaviors.

### Ultrasonic vocalization analysis

The recorded ultrasonic vocalizations (USVs) were analyzed using a custom-written software and visually corrected by experimenters. Analysis of the USV emitter was performed by comparing sound levels of each USV in the four microphones using custom-written MATLAB codes and combining this with the top view video of the experiment to visually identify the emitter, based on the rats’ head direction and proximity to the microphones. Emitter of each USV was labelled as either ‘observer, ‘playback’, ‘demonstrator, or ‘unclear’ (particularly in case the USVs were emitted while snouts of both rats were touching).

### Electrophysiology

Single and multi unit activity in the left trunk somatosensory cortex (1 mm posterior, 2-4 mm lateral from Bregma) of the observer animal was recorded using a Harlan 8 drive, Cheetah software, and DigitalLynx SX (Neuralynx, Bozeman, MT, USA) and analyzed as previously reported [59]. Units with average firing rate < 0.1 Hz were excluded. Unless otherwise denoted, response index of a given phase was calculated for each unit as RI_phase_ = (FR_phase_ – FR_pre_) / (FR_phase_ + FR_pre_), where FR_phase_ is the mean firing rate during a phase of interest, and FR_pre_ is the mean firing rate from −5 to −1 s period relative to the onset of the phase of interest. Units with RI_witnessed_dorsal_ > 0.1 and RI_direct_dorsal_ > 0.1 were defined as tickling mirror units (Figure 2B), which is stricter criteria than those used in a previous study of mirror neurons [21]. Unless otherwise denoted, peristimulus time histograms (PSTHs) were quantified by comparing firing rate during [−5, 0] s and [0, 5] s to the onset of a given phase using signed-rank test.

### Histology

After the last experiment, animals were anaesthetized and the tetrode tracks were labeled with electrolytic lesions by applying a DC current (8 s, 8 µA, electrode tip negative). Animals then received an overdose of isoflurane and were transcardially perfused with a pre-fixative solution followed by a 4% paraformaldehyde (PFA) solution. For histological processing brains were cut in 100 µm coronal sections and stained for cytochrome-oxidase activity as described previously [60]. Assignment of recording sites to layers was done based on the layer-specific staining.

### Statistical analysis

Normally distributed data (Shapiro-Wilk test) are shown as mean ± SEM and intergroup comparisons were performed with paired-*t*-test for paired data, and *t*-test for non-paired data. Data with non-normal distribution were evaluated with signed-rank test for paired data and rank-sum test for non-paired data. *n* refers to sample size. Data were analyzed using MATLAB 2017b and 2018b, and Python 2.7 and 3.6.

**Figure S1.**
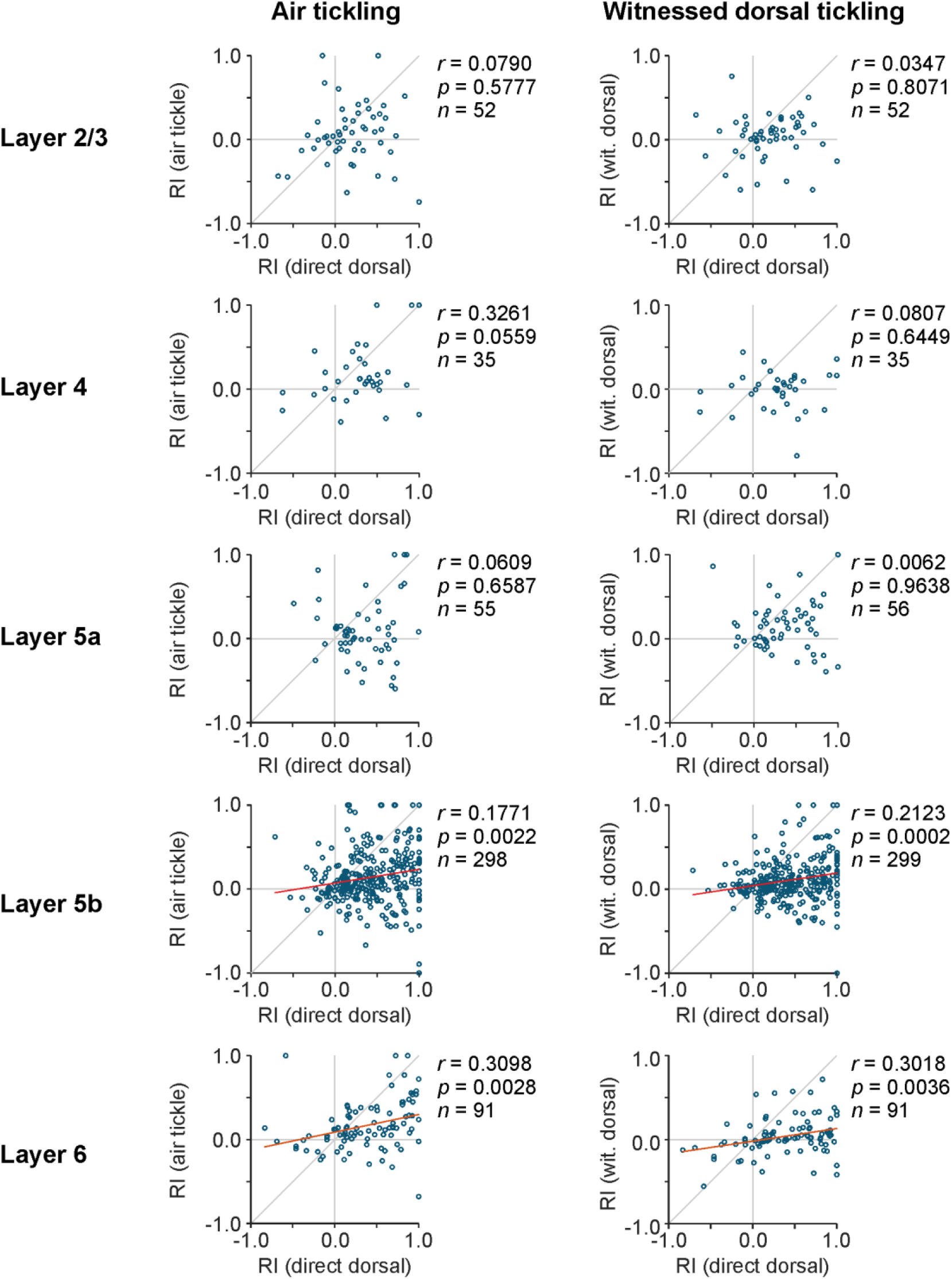
Tickle responses predict air tickling and witnessed tickle responses in deep-layer trunk cortex. Left, scatter plots show response indices (RI) for dorsal tickling (dorsal) vs. air tickle in layers 2/3, 4, 5a, 5b and 6. Grey diagonal lines are unity line. Red lines are linear fit in case of significant correlation (Pearson’s). Right, same as left but for witnessed dorsal tickling of demonstrator (wit. dorsal). *n*: number of units.

**Figure S2.**
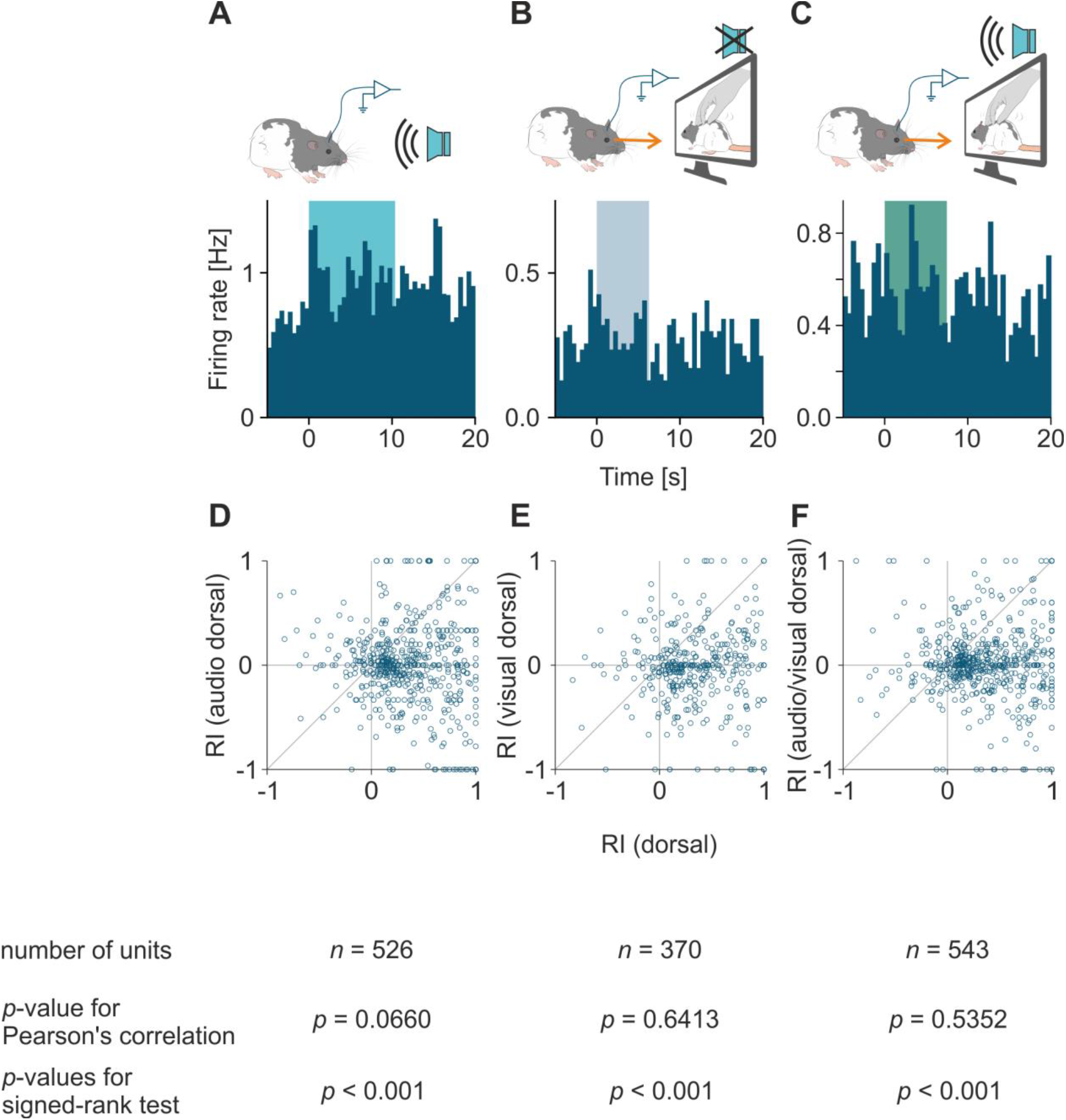
Neuronal response to playback of tickling footage. A. Top, schematic illustrations of neuronal recording during audio playback of dorsal tickling footage. Bottom, population peri-stimulus time histograms of firing rate. Data are binned to 0.5 s. With of the color boxes indicate median duration of events. B. Same as A but for visual playback of dorsal tickling. C. Same as A but for audio/visual playback of dorsal tickling. D. Scatter plot shows response index of dorsal tickling vs. audio playback of dorsal tickling (audio dorsal). Number of units, *p*-values for Pearson’s correlation, *p*-values for signed-rank test are described below. E. Same as D but for dorsal tickling vs. visual playback of dorsal tickling. F.Same as D but for dorsal tickling vs. audio/visual playback of dorsal tickling.

**Figure S3.**
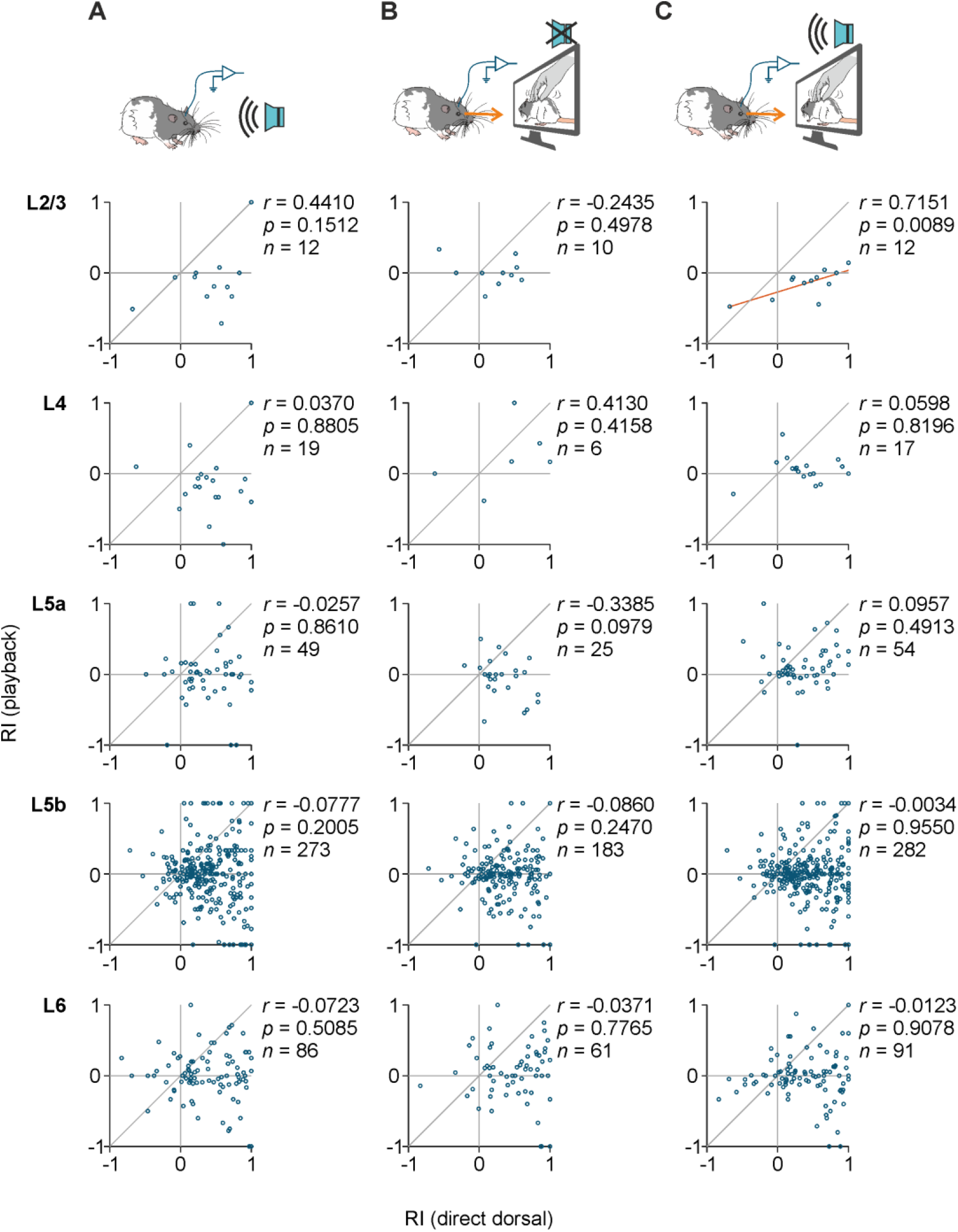
No systematic correlations between tickle response and witnessed tickle response across layers. A. Top, schematic illustrations of neuronal recording during audio playback of dorsal tickling footage. Bottom, scatter plots show response index of dorsal tickling vs. audio playback of dorsal tickling in each cortical layer. *r*: Pearson’s correlation coefficient; *p*: *p*-values for Pearson’s correlation; *n*: number of units. B. Same as A but for visual playback. C. Same as A but for audio/visual playback.

**Figure S4.**
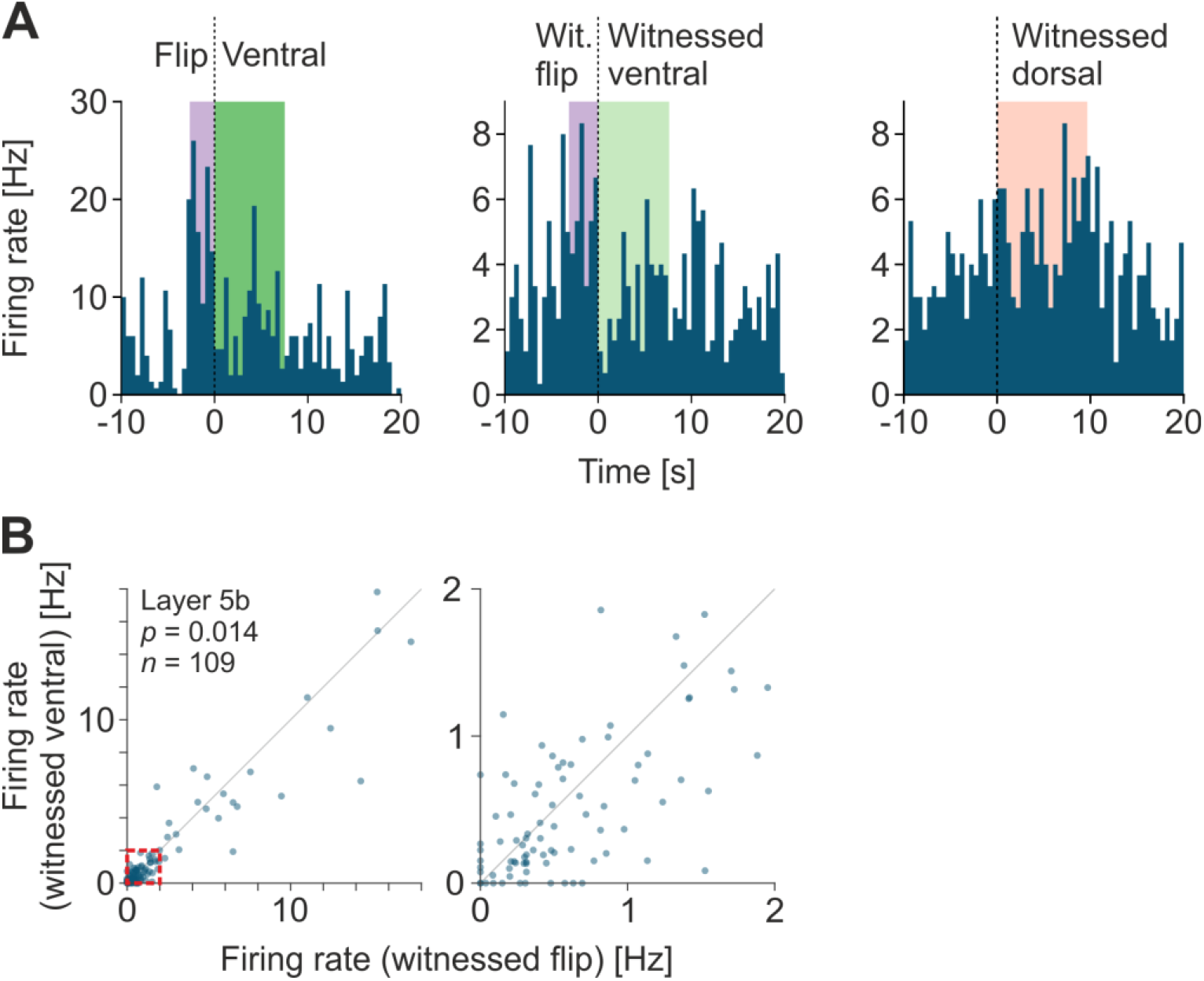
Action specificity of neuronal response to witnessed tickling. A. Peri-stimulus time histograms of firing rate of a representative unit aligned to observer ventral tickling (left, following flip), witnessed ventral tickling (middle), and witnessed dorsal tickling (right). Note the robust response to flip as well as witnessed flip, and rather suppressed firing rate in the following ventral and witnessed ventral tickling. Data are binned to 0.5 s. Width of color boxes indicate median duration of events. B. Firing rate during witnessed flip vs. during witnessed ventral tickling of layer 5b trunk cortex neurons (*n* = 109 mirror units). Grey line is unity line. Data in the red box is shown in the right axes. The *p*-value indicates signed-rank test.

**Movie S1**

The observer rat witnesses the demonstrator rat being tickled. The observer rat exhibits Freudensprünge, suggesting contagion of tickling-evoked positive emotions.

## References

1. Panksepp, J., and Panksepp, J.B. (2013). Toward a cross-species understanding of empathy. Trends Neurosci. 36, 489–496.

2. Hatfield, E., Cacioppo, J.T., and Rapson, R.L. (1993). Emotional Contagion. Current Directions in Psychological Science.

3. Darwin, C. (1871). The descent of man : and selection in relation to sex, (London: J. Murray).

4. Nakahashi, W., and Ohtsuki, H. (2015). When is emotional contagion adaptive? J. Theor. Biol. 380, 480–488.

5. de Waal, F.B.M., and Preston, S.D. (2017). Mammalian empathy: behavioural manifestations and neural basis. Nature Reviews Neuroscience 18, 498-+.

6. Paradiso, E., Gazzola, V., and Keysers, C. (2021). Neural mechanisms necessary for empathy-related phenomena across species. Curr. Opin. Neurobiol. 68, 107–115.

7. de Waal, F.B.M. (2008). Putting the Altruism Back into Altruism: The Evolution of Empathy. Annual Review of Psychology.

8. Meyza, K.Z., Bartal, I.B.A., Monfils, M.H., Panksepp, J.B., and Knapska, E. (2017). The roots of empathy: Through the lens of rodent models.

9. Rizzolatti, G., Fogassi, L., and Gallese, V. (2001). Neurophysiological mechanisms underlying the understanding and imitation of action. Nat. Rev. Neurosci. 2, 661–670.

10. Di Pellegrino, G., Fadiga, L., Fogassi, L., Gallese, V., and Rizzolatti, G. (1992). Understanding motor events: a neurophysiological study. Exp. Brain Res. 91, 176–180.

11. Gallese, V., Fadiga, L., Fogassi, L., and Rizzolatti, G. (1996). Action recognition in the premotor cortex. Brain 119, 593–609.

12. Fogassi, L., Ferrari, P.F., Gesierich, B., Rozzi, S., Chersi, F., and Rizzolatti, G. (2005). Parietal lobe: from action organization to intention understanding. Science 308, 662–667.

13. Shepherd, S.V., Klein, J.T., Deaner, R.O., and Platt, M.L. (2009). Mirroring of attention by neurons in macaque parietal cortex. Proceedings of the National Academy of Sciences 106, 9489–9494.

14. Tkach, D., Reimer, J., and Hatsopoulos, N.G. (2007). Congruent activity during action and action observation in motor cortex. J. Neurosci. 27, 13241–13250.

15. Ishida, H., Nakajima, K., Inase, M., and Murata, A. (2010). Shared mapping of own and others’ bodies in visuotactile bimodal area of monkey parietal cortex. J. Cognit. Neurosci. 22, 83–96.

16. Fujii, N., Hihara, S., and Iriki, A. (2007). Dynamic social adaptation of motion-related neurons in primate parietal cortex. PloS One 2, e397.

17. Breveglieri, R., Vaccari, F.E., Bosco, A., Gamberini, M., Fattori, P., and Galletti, C. (2019). Neurons modulated by action execution and observation in the macaque medial parietal cortex. Curr. Biol. 29, 1218-1225. e1213.

18. Farina, E., Borgnis, F., and Pozzo, T. (2020). Mirror neurons and their relationship with neurodegenerative disorders. J. Neurosci. Res. 98, 1070–1094.

19. Smith, M.L., Asada, N., and Malenka, R.C. (2021). Anterior cingulate inputs to nucleus accumbens control the social transfer of pain and analgesia. 371, 153–159.

20. Yu, Y.Q., Barry, D.M., Hao, Y., Liu, X.T., and Chen, Z.F. (2017). Molecular and neural basis of contagious itch behavior in mice. Science 355, 1072–1076.

21. Carrillo, M., Han, Y., Migliorati, F., Liu, M., Gazzola, V., and Keysers, C. (2019). Emotional Mirror Neurons in the Rat’s Anterior Cingulate Cortex. Curr. Biol.

22. Adolphs, R., Damasio, H., Tranel, D., Cooper, G., and Damasio, A.R. (2000). A role for somatosensory cortices in the visual recognition of emotion as revealed by three-dimensional lesion mapping. J. Neurosci.

23. Kragel, P.A., and LaBar, K.S. (2016). Somatosensory Representations Link the Perception of Emotional Expressions and Sensory Experience. eNeuro 3.

24. Keysers, C., Kaas, J.H., and Gazzola, V. (2010). Somatosensation in social perception. Nature Reviews Neuroscience 11, 417–428.

25. Damasio, A.R. (1996). The somatic marker hypothesis and the possible functions of the prefrontal cortex. Philos T R Soc B 351, 1413–1420.

26. Provine, R.R. (2004). Laughing, tickling, and the evolution of speech and self. Current Directions in Psychological Science 13, 215–218.

27. Panksepp, J., and Burgdorf, J. (1999). Laughing rats? Playful tickling arouses high frequency ultrasonic chirping in young rodents. In Toward a Science of Consciousness III, S.R. Hameroff, D. Chalmers and A.W. Kaszniak, eds. (MIT Press), pp. 231–244.

28. Ishiyama, S., and Brecht, M. (2016). Neural correlates of ticklishness in the rat somatosensory cortex. Science 354, 757–760.

29. Ishiyama, S., Kaufmann, L.V., and Brecht, M. (2019). Behavioral and Cortical Correlates of Self-Suppression, Anticipation, and Ambivalence in Rat Tickling. Curr. Biol. 29, 3153–3164.

30. Knutson, B., Burgdorf, J., and Panksepp, J. (2002). Ultrasonic Vocalizations as indices of affective states in rats. Psychol. Bull. 128, 961–977.

31. Reinhold, A.S., Sanguinetti-Scheck, J.I., Hartmann, K., and Brecht, M. (2019). Behavioral and neural correlates of hide-and-seek in rats. Science 365, 1180–1183.

32. Osvath, M., and Sima, M. (2014). Sub-adult ravens synchronize their play: a case of emotional contagion. Animal Behavior and Cognition 1, 197–205.

33. Schwing, R., Nelson, X.J., Wein, A., and Parsons, S. (2017). Positive emotional contagion in a New Zealand parrot. Curr. Biol. 27, R213–R214.

34. Provine, R.R. (1992). Contagious laughter: Laughter is a sufficient stimulus for laughs and smiles. Bulletin of the Psychonomic Society.

35. Davila-Ross, M., Allcock, B., Thomas, C., and Bard, K.A. (2011). Aping expressions? Chimpanzees produce distinct laugh types when responding to laughter of others. Emotion 11, 1013.

36. Pellis, S.M., and McKenna, M.M. (1992). Intrinsic and extrinsic influences on play fighting in rats: effects of dominance, partner’s playfulness, temperament and neonatal exposure to testosterone propionate. Behav. Brain Res. 50, 135–145.

37. Varlinskaya, E.I., Spear, L.P., and Spear, N.E. (1999). Social behavior and social motivation in adolescent rats: role of housing conditions and partner’s activity. Physiol. Behav. 67, 475–482.

38. Hammond, T., Bombail, V., Nielsen, B.L., Meddle, S.L., Lawrence, A.B., and Brown, S.M. (2019). Relationships between play and responses to tickling in male juvenile rats. Appl. Anim. Behav. Sci. 221.

39. Kisko, T.M., Euston, D.R., and Pellis, S.M. (2015). Are 50-khz calls used as play signals in the playful interactions of rats? III. The effects of devocalization on play with unfamiliar partners as juveniles and as adults. Behav. Processes 113, 113–121.

40. Bierley, R.A., Hughes, S.L., and Beatty, W.W. (1986). Blindness and play fighting in juvenile rats. Physiol. Behav. 36, 199–201.

41. Siviy, S.M., and Panksepp, J. (1987). Sensory modulation of juvenile play in rats. Dev. Psychobiol. 20, 39–55.

42. Brecht, M. (2017). The Body Model Theory of Somatosensory Cortex. Neuron 94, 985–992.

43. Kohler, E., Keysers, C., Umilta, M.A., Fogassi, L., Gallese, V., and Rizzolatti, G. (2002). Hearing sounds, understanding actions: action representation in mirror neurons. Science 297, 846–848.

44. Keysers, C., Kohler, E., Umiltà, M.A., Nanetti, L., Fogassi, L., and Gallese, V. (2003). Audiovisual mirror neurons and action recognition. Exp. Brain Res. 153, 628–636.

45. Gazzola, V., Aziz-Zadeh, L., and Keysers, C. (2006). Empathy and the somatotopic auditory mirror system in humans. Curr. Biol. 16, 1824–1829.

46. Gharaei, S., Honnuraiah, S., Arabzadeh, E., and Stuart, G.J. (2020). Superior colliculus modulates cortical coding of somatosensory information. Nature communications 11, 1–14.

47. Burn, C.C. (2008). What is it like to be a rat? Rat sensory perception and its implications for experimental design and rat welfare. Appl. Anim. Behav. Sci. 112, 1–32.

48. Vielle, C., Montanari, C., Pelloux, Y., and Baunez, C. (2021). Evidence for a vocal signature in the rat and its reinforcing effects. bioRxiv, 2021.2006.2007.447373.

49. Gonzalez-Liencres, C., Juckel, G., Tas, C., Friebe, A., and Brune, M. (2014). Emotional contagion in mice: the role of familiarity. Behav. Brain Res. 263, 16–21.

50. Rogers-Carter, M.M., Djerdjaj, A., Culp, A.R., Elbaz, J.A., and Christianson, J.P. (2018). Familiarity modulates social approach toward stressed conspecifics in female rats. PloS One 13, e0200971.

51. Cox, S.S., and Reichel, C.M. (2020). Rats display empathic behavior independent of the opportunity for social interaction. Neuropsychopharmacology 45, 1097–1104.

52. Langford, D.J., Crager, S.E., Shehzad, Z., Smith, S.B., Sotocinal, S.G., Levenstadt, J.S., Chanda, M.L., Levitin, D.J., and Mogil, J.S. (2006). Social modulation of pain as evidence for empathy in mice. Science.

53. Andraka, K., Kondrakiewicz, K., Rojek-Sito, K., Ziegart-Sadowska, K., Meyza, K., Nikolaev, T., Hamed, A., Kursa, M., Wojcik, M., Danielewski, K., et al. (2021). Distinct circuits in rat central amygdala for defensive behaviors evoked by socially signaled imminent versus remote danger. Curr. Biol. 31, 2347–2358 e2346.

54. Keysers, C., and Gazzola, V. (2021). Emotional contagion: Improving survival by preparing for socially sensed threats. Curr. Biol. 31, R728–R730.

55. Pitcher, D., Garrido, L., Walsh, V., and Duchaine, B.C. (2008). Transcranial magnetic stimulation disrupts the perception and embodiment of facial expressions. J. Neurosci. 28, 8929–8933.

56. Adolphs, R. (2002). Neural systems for recognizing emotion. Curr. Opin. Neurobiol. 12, 169–177.

57. Banissy, M.J., Sauter, D.A., Ward, J., Warren, J.E., Walsh, V., and Scott, S.K. (2010). Suppressing sensorimotor activity modulates the discrimination of auditory emotions but not speaker identity. J. Neurosci. 30, 13552–13557.

58. Lausberg, H., and Sloetjes, H. (2009). Coding gestural behavior with the NEUROGES-ELAN system. Behav. Res. Methods 41, 841–849.

59. Rao, R.P., Mielke, F., Bobrov, E., and Brecht, M. (2014). Vocalization-whisking coordination and multisensory integration of social signals in rat auditory cortex. eLife 3, e03185.

60. Brecht, M., and Sakmann, B. (2002). Dynamic representation of whisker deflection by synaptic potentials in spiny stellate and pyramidal cells in the barrels and septa of layer 4 rat somatosensory cortex. J. Physiol. 543, 49–70.

